# Paradoxical activation of a type VI secretion system (T6SS) phospholipase effector by its cognate immunity protein

**DOI:** 10.1101/2023.03.28.534661

**Authors:** Steven J. Jensen, Zachary C. Ruhe, August F. Williams, Dinh Q. Nhan, Fernando Garza-Sánchez, David A. Low, Christopher S. Hayes

**Affiliations:** Department of Molecular, Cellular and Developmental Biology, University of California, Santa Barbara, Santa Barbara, CA 93106-9625, USA; Biomolecular Science and Engineering Program, University of California, University of California, Santa Barbara, Santa Barbara, CA 93106-9625, USA

## Abstract

Type VI secretion systems (T6SS) deliver cytotoxic effector proteins into target bacteria and eukaryotic host cells. Antibacterial effectors are invariably encoded with cognate immunity proteins that protect the producing cell from self-intoxication. Here, we identify transposon insertions that disrupt the *tli* immunity gene of *Enterobacter cloacae* and induce auto-permeabilization through unopposed activity of the Tle phospholipase effector. This hyper-permeability phenotype is T6SS-dependent, indicating that the mutants are intoxicated by Tle delivered from neighboring sibling cells rather than by internally produced phospholipase. Unexpectedly, an in-frame deletion of *tli* does not induce hyper-permeability because Δ*tli* null mutants fail to deploy active Tle. Instead, the most striking phenotypes are associated with disruption of the *tli* lipoprotein signal sequence, which prevents immunity protein localization to the periplasm. Immunoblotting reveals that most hyper-permeable mutants still produce Tli, presumably from alternative translation initiation codons downstream of the signal sequence. These observations suggest that cytosolic Tli is required for the activation and/or export of Tle. We show that Tle growth inhibition activity remains Tli-dependent when phospholipase delivery into target bacteria is ensured through fusion to the VgrG β-spike protein. Together, these findings indicate that Tli has distinct functions depending on its subcellular localization. Periplasmic Tli acts as a canonical immunity factor to neutralize incoming effector proteins, while a cytosolic pool of Tli is required to activate the phospholipase domain of Tle prior to T6SS-dependent export.

## Importance

Gram-negative bacteria use type VI secretion systems deliver toxic effector proteins directly into neighboring competitors. Secreting cells also produce specific immunity proteins that neutralize effector activities to prevent auto-intoxication. Here, we show that the Tli immunity protein of *Enterobacter cloacae* has two distinct functions depending on its subcellular localization. Periplasmic Tli acts as a canonical immunity factor to block Tle lipase effector activity, while cytoplasmic Tli is required to activate the lipase prior to export. These results indicate that Tle interacts transiently with its cognate immunity protein to promote effector protein folding and/or packaging into the secretion apparatus.

## Introduction

Research over the past twenty years shows that bacteria commonly deliver growth inhibitory toxins directly into neighboring competitors (1). This phenomenon was first described as contact-dependent growth inhibition or “CDI” in *Escherichia coli* isolate EC93 (2). CDI is mediated by a specialized type V secretion system (T5SS) composed of CdiB and CdiA proteins. CdiB is an Omp85 family β-barrel protein responsible for transport of the CdiA effector protein to the cell surface (3). Upon binding a receptor on a neighboring bacterium, CdiA delivers its C-terminal toxin domain into the target cell to inhibit growth. CDI^+^ strains protect themselves from auto-inhibition by producing immunity proteins that specifically neutralize the toxin domain. Shortly after the discovery of T5SS-mediated CDI, two reports demonstrated that type VI secretion systems (T6SS) are also potent mediators of interbacterial competition (4, 5). T6SSs are dynamic multiprotein machines related in structure and function to the contractile tails of *Myoviridae* bacteriophages. The T6SS duty cycle begins with the assembly of a phage-like baseplate complex around trimeric VgrG, which is homologous to the β-tailspike protein of coliphage T4 (6). The baseplate then docks onto the cytosolic face of the transenvelope complex, which forms a secretion conduit across the cell envelope (7). This latter assembly nucleates the polymerization of a contractile sheath surrounding a central tube of Hcp proteins. Toxic effector proteins are typically linked to the VgrG β-spike (8, 9), and can also be packaged within the lumen of the Hcp tube (10). Once the sheath has grown to span the width of the cell, it contracts to expel the VgrG-capped tube through the transenvelope complex. The ejected projectile perforates neighboring bacteria and releases its toxic payload to inhibit target cell growth. As with toxic CdiA proteins, antibacterial T6SS effectors are encoded with cognate immunity proteins that protect the cell from intoxication by its isogenic siblings (1, 11). In this manner, T5S and T6S systems confer a competitive fitness advantage to bacteria, and their effector-immunity protein pairs play an important role in self/nonself discrimination.

Given that T5SS-and T6SS-mediated competition relies on cell-to-cell contact, the systems are often only deployed under conditions that favor productive effector delivery. Several regulatory strategies have been described for T6SSs, which are controlled at all levels of gene expression. T6SS loci are commonly subject to H-NS mediated transcriptional silencing (12-16), and many require specific bacterial enhancer proteins to initiate transcription in conjunction with 0^54^ (17, 18). Environmental signals also influence T6SS transcription through quorum sensing and two-component regulatory signaling pathways (16, 19, 20). In *Pseudomonas aeruginosa*, the stability and translation of T6SS transcripts is modulated by small non-coding RNAs (16). Finally, assembly of the secretion apparatus is subject to post-translational control by a serine/threonine kinase and phosphatase (21).

There are fewer studies on the regulation of antibacterial T5SSs. The originally identified *cdiBAI* locus from *E. coli* EC93 is expressed constitutively (2, 22), but systems from other species are clearly regulated. The CDI system of *Burkholderia thailandensis* E264 is activated through a quorum sensing pathway (23), and the loci of *Dickeya dadantii* EC16 and 3937 are only expressed when these bacteria are grown on plant hosts (24-26). The latter observations suggest that CDI promotes host colonization and pathogenesis, though the T5SS effector domains deployed by these phytopathogens are clearly antibacterial.

We previously reported that the CDI system of *Enterobacter cloacae* ATCC 13047 (ECL) is not active under laboratory conditions, though it is functional when expressed from a heterologous promoter (27). The wild-type *cdiBAI*^ECL^ operon appears to be transcribed from a single promoter with possible operator sites for PurR and Fnr. These predictions suggest that *cdi* transcription may be repressed when ECL is grown in standard culture media. In an attempt to identify transcriptional regulators, we generated an ECL strain that harbors the *gusA* reporter gene under the control of the *cdi* promoter, then screened for transposon mutants that exhibit increased β-glucuronidase activity.

Surprisingly, the recovered mutations have no effect on *cdi* transcription, and instead act to increase cell permeability to the chromogenic β-glucuronidase substrate. We identified multiple insertions in *tli*, which encodes a previously described immunity protein that neutralizes the <u>T</u>6SS phospholipase <u>e</u>ffector Tle (28). Thus, cell envelope integrity is disrupted by unrestrained phospholipase activity in these mutants. Increased permeability is T6SS-dependent, indicating that *tli* mutants are intoxicated by Tle delivered from neighboring siblings rather than internally produced phospholipase. Notably, we find that Δ*tli* null mutants are not hyper-permeable because they fail to deploy active Tle. Taken together, our findings indicate that Tli has distinct functions depending on its subcellular localization. In addition to its established role as a periplasmic immunity factor, we propose that a cytosolic pool of Tli is required to activate the Tle phospholipase prior to T6SS-dependent export.

## Results

### Disruption of tli increases cell permeability

We initially sought to identify regulators of the *cdiBAI* gene cluster in *Enterobacter cloacae* ATCC 13047 (ECL) using a β-glucuronidase (*gusA*) reporter to monitor expression. We fused the *E. coli gusA* gene to the *cdiB* (ECL_04452) promoter region and placed the construct at the ECL *glmS* locus using Tn*7*-mediated transposition. The resulting *cdi-gusA* reporter strain was subjected to transposon Tn*5*-mediated mutagenesis and >25,000 insertion mutants were screened on agar medium supplemented with the chromogenic β-glucuronidase substrate, X-gluc. Nine different transposon insertion sites were identified from 17 mutants that formed pigmented colonies on indicator medium.

One insertion is located within *tle* (ECL_01553), and the other eight disrupt the downstream *tli* cistron (ECL_01554) (**Fig. 1A**). The *tle* gene encodes a toxic phospholipase effector deployed by the type VI secretion system-1 (T6SS-1) of ECL, and *tli* encodes an ankyrin-repeat immunity protein that neutralizes Tle activity (29). We transferred the transposon mutations into the original parental reporter strain using Red-mediated recombineering and found that all of the reconstructed mutants, except *tle-1106*, recapitulate the original phenotype on indicator medium (**Fig. 1B**). Quantification of β-glucuronidase activity in cell lysates from a subset of mutants revealed that each has the same specific activity as the parental reporter strain (**Fig. 1C**). These data indicate that *cdi* transcription is unaffected by the *tli* mutations. Moreover, immunoblot analyses showed that none of the mutants exhibit increased production of CdiB (**Fig. 1D**) or CdiA (**Fig. 1E**). Given these results, we hypothesized that the loss of Tli immunity function leads to unopposed Tle phospholipase activity, which in turn degrades cell membranes to increase permeability to X-gluc. Consistent with this model, parental reporter cells grown on indicator medium become pigmented after treatment with SDS solution (**Fig. 2A**). Moreover, complementation with plasmid-borne *tli* suppresses pigment conversion (**Fig. 2B**), but has little effect on specific β-glucuronidase activities (**Fig. 2C**). Together, these results indicate that diminished immunity function leads to increased cell permeability.

**Fig. 1.**
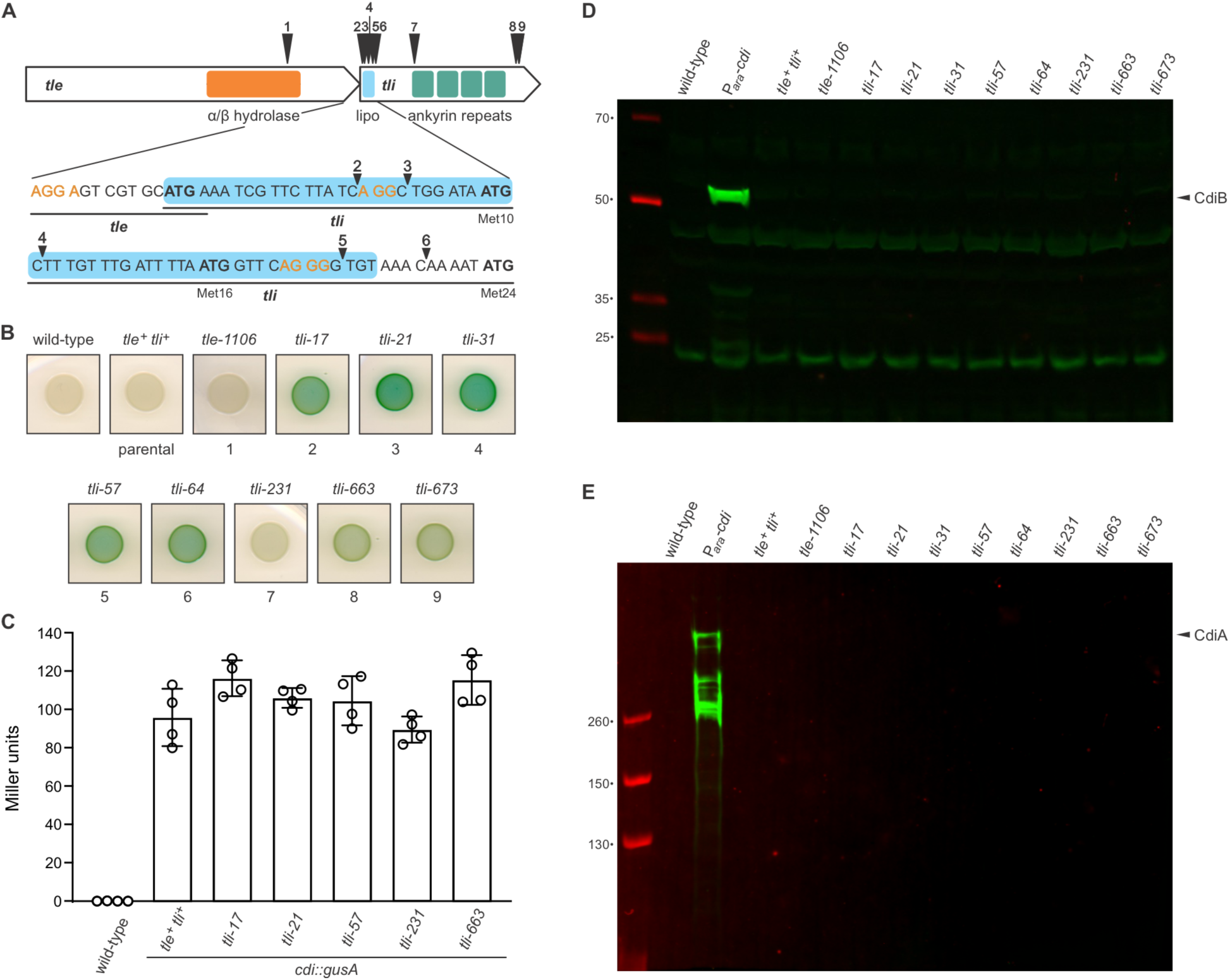
Identification of Tn*5* insertions in the *tli* immunity gene. **A**) Schematic of Tn*5* insertion sites in *tle* and *tli*. Regions encoding the predicted α/β-hydrolase domain of Tle and the Tli lipoprotein signal peptide (lipo) and ankyrin repeats are depicted. Alternative translation initiation codons (Met10, Met16 and Met24) are indicated, and possible ribosome-binding sites are shown in orange font. **B**) Wild-type ECL, the parental *cdi-gusA* reporter strain (*tle^+^ tli^+^*) and *tli* mutants were grown on LB-agar supplemented with X-gluc. **C**) Quantification of β-glucuronidase activity in selected *tli* mutants. Activities were quantified as Miller units as described in the Methods. Data for technical replicates from two independent experiments are shown with the average ± standard deviation. **D** & **E**) Tn*5* insertions do not upregulate CdiB or CdiA production. Protein samples from wild-type ECL, the parental *tle^+^ tli^+^* reporter strain and *tli* mutants were subjected to immunoblot analysis using polyclonal antisera to CdiB (**D**) and CdiA (**E**). The P*_ara_*-*cdi* sample is from an ECL strain that contains an arabinose-inducible promoter inserted upstream of the native *cdiBAI* gene cluster.

**Fig. 2.**
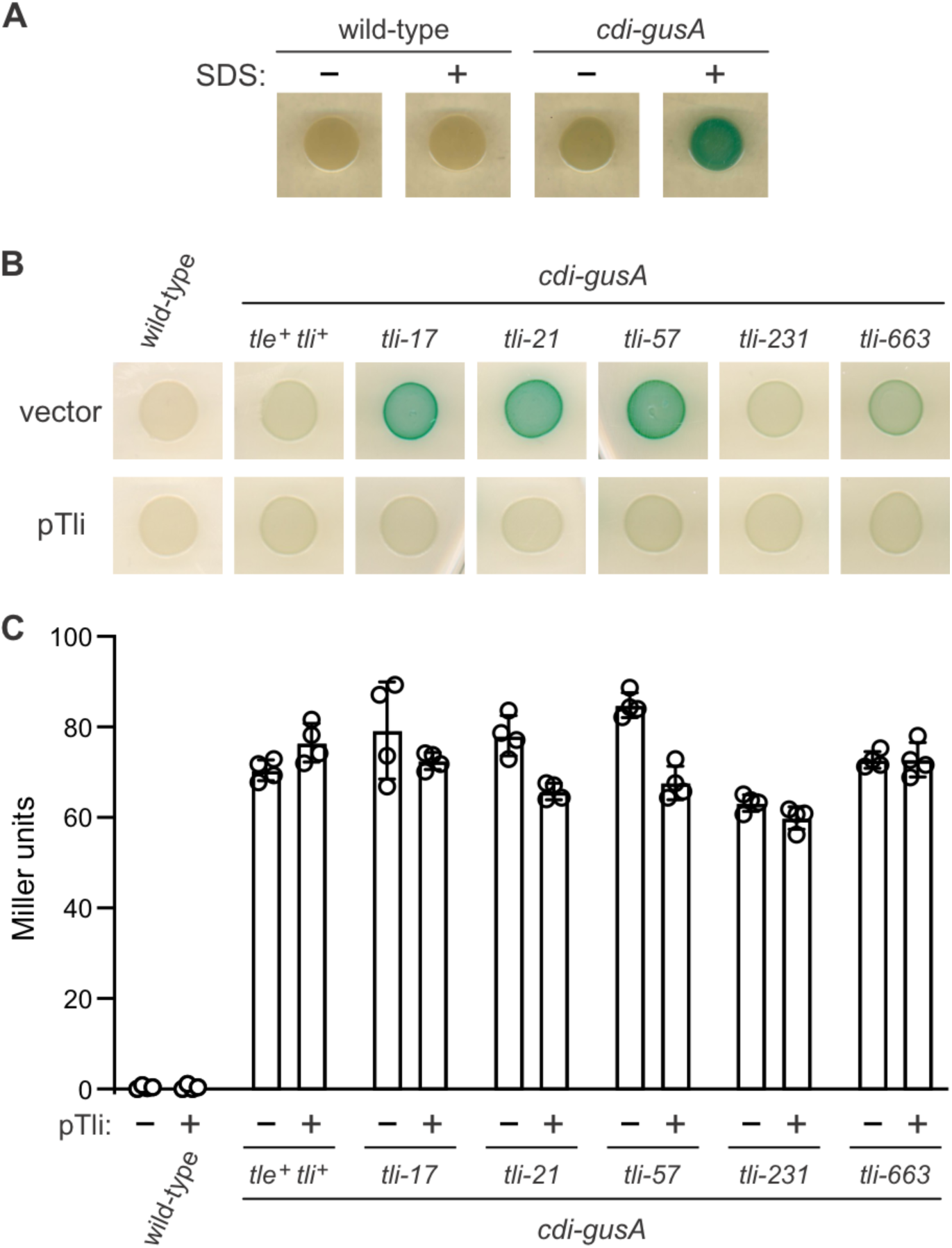
Disruption of *tli* increases cell permeability. **A**) ECL is impermeable to chromogenic X-gluc substrate. Wild-type and *cdi-gusA* reporter cells were grown on LB-agar supplemented with X-gluc. Where indicated, SDS solution (1% wt/vol) was applied to the edge of the colonies prior to imaging. **B**) The indicated ECL strains were complemented with plasmid-borne *tli* (pTli) or an empty vector and grown on LB-agar supplemented with X-gluc. **C**) Complementation with *tli* does not affect *cdi-gusA* reporter expression. β-glucuronidase activities were quantified as Miller units as described in the Methods. Data for technical replicates from two independent experiments are shown with the average ± standard deviation.

### Increased cell permeability is dependent on T6SS-1 activity

Two scenarios could account for hyper-permeability: the *tli* mutants could be permeabilized by internally produced phospholipase, or they could be intoxicated by Tle delivered from neighboring sibling cells. These models are not exclusive and the phenotype could reflect a combination of external and internal phospholipase activity. To eliminate the contribution from T6SS-delivered effector, we deleted *tssM* (**Fig. 3A**), which encodes a critical component of the transenvelope complex required for secretion. The resulting *tli ΔtssM* cells do not secrete Hcp into culture supernatants (**Fig. 3B**), confirming that T6SS-1 is inactivated in these strains. The *tli* Δ*tssM* strains also exhibit less pigmentation when grown on X-gluc indicator medium (**Fig. 3C**, top row). To ensure that the Δ*tssM* mutation does not abrogate *tle* expression, we showed that complementation with plasmid-borne *tssM* restores Hcp secretion and hyper-permeability phenotypes to each strain (**Figs. 3B** & **3C**, bottom row). These data indicate that increased cell permeability is largely due to T6SS-1 dependent exchange of phospholipase toxin between sibling cells.

**Fig. 3.**
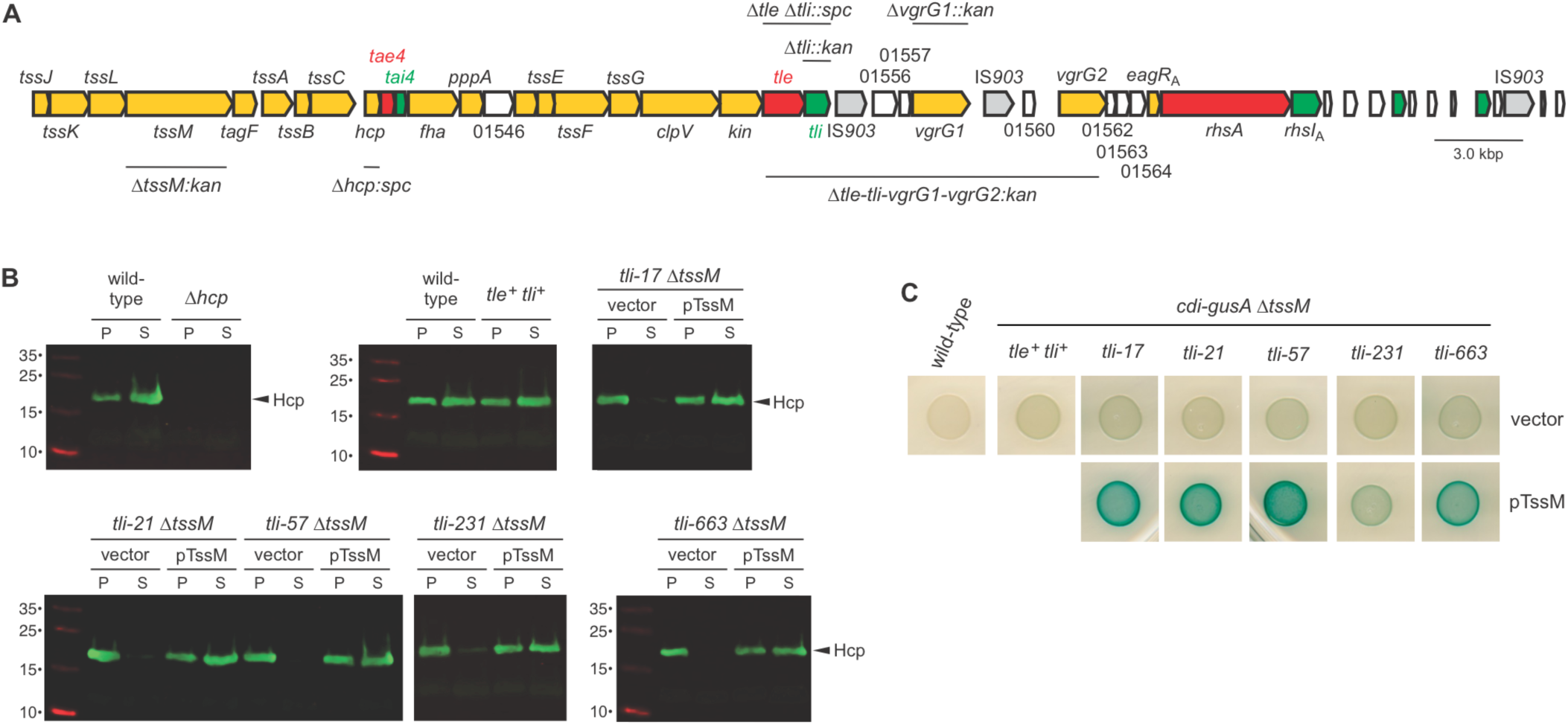
Hyper-permeability is T6SS-1 dependent. **A**) Schematic of the ECL T6SS-1 locus. Core secretion system genes are colored light orange, and effector and immunity genes are shown in red and green, respectively. Genes of unknown function are not colored and are indicated by their ordered locus number. The extent of the gene deletions used in this study are indicated. **B**) ECL Δ*tssM* strains do not secrete Hcp. The indicated ECL strains were grown in LB, then cell pellet (P) and culture supernatant (S) fractions prepared as described in Methods. Where indicated, the strains carried a plasmid-borne copy of *tssM* (pTssM) or an empty vector plasmid. Fractions were analyzed by SDS-PAGE and immunoblotting with polyclonal antisera to Hcp. **C**) *tli* Δ*tssM* mutants do not exhibit increased cell permeability. The indicated ECL strains were grown on LB-agar supplemented with X-gluc. Where indicated, the strains carried a plasmid-borne copy of *tssM* (pTssM) or an empty vector plasmid.

Although *tli* is disrupted in each mutant, most retain the capacity to produce a truncated form of the immunity protein. Insertions in the lipoprotein signal sequence (*tli-17, tli-21, tli-31, tli-57* and *tli-64*) should allow cytosolic Tli to be synthesized using Met10, Met16 or Met24 as alternative translation initiation codons (**Fig. 1A**). Further, the *tli-663* and *tli-673* alleles are predicted to produce secreted Tli lipoproteins that carry transposon-encoded residues fused at Asp221 and Lys224, respectively (**Fig. 1A**). Immunoblotting revealed that the *tli-17, tli-21, tli-57* and *tli-64* mutants accumulate Tli to about the same level as the parental *tli^+^* strain, but we were unable to detect antigen in *tli-231*, *tli-663* and *tli-763* mutant lysates (**Fig. 4A**). We next asked whether truncated Tli provides any protection against Tle effector delivered from wild-type ECL inhibitor cells. Because the *tli* mutants also deploy Tle to intoxicate their siblings, we examined immunity function in a Δ*tssM* background to ensure that the mutants act only as target cells during co-culture. The *tli ΔtssM* target strains are outcompeted between 50-and 100-fold by wild-type ECL, but they show no fitness disadvantage when co-cultured with ECL Δ*tssM* mock inhibitor cells (**Fig. 4B**). This level of growth inhibition is comparable to that observed with *Δtle Δtli* target cells (**Fig. 4B**), which harbor a true null allele of *tli*. Thus, the transposon insertions completely abrogate immunity to Tle. Given that cytosolic immunity protein fails to protect, these results also indicate that Tli must be localized to the periplasm to neutralize phospholipase activity.

**Fig. 4.**
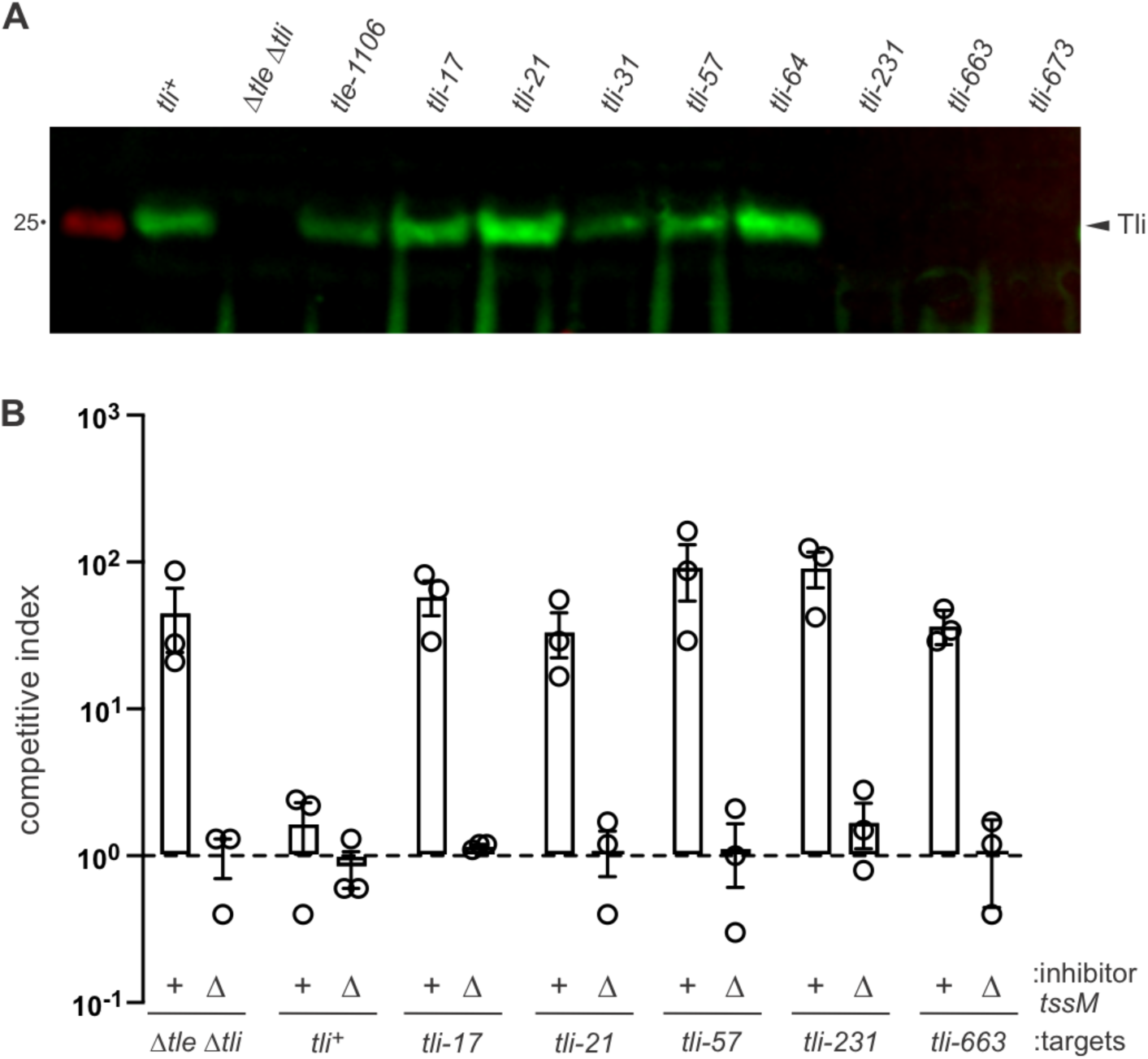
*tli* mutants produce Tli, but are not immune to Tle effector. **A**) *tli* mutants carrying Tn*5* insertions in the lipoprotein signal sequence still produce immunity protein. Protein samples from the indicated ECL strains were subjected to immunoblot analysis using polyclonal antisera to Tli. **B**) Tn*5* insertions ablate the immunity function of Tli. Wild-type ECL (*tssM^+^*) and mock (Δ*tssM*) inhibitor strains were mixed with the indicated *tli ΔtssM* target cells and spotted onto LB-agar. After 4 h of co-culture, inhibitor and target cells were enumerated as colony forming units (cfu) on selective media. The competitive index equals the final ratio of inhibitor to target cells divided by the initial ratio. Presented data are the average ± standard error from three independent experiments.

### Tli is required to deploy Tle

Although the *tli* disruptions behave as null mutations in competition co-cultures, we noted that the *tli-231* insertion induces a modest permeability phenotype compared to the other alleles. This observation is curious because *tli-231* is arguably the most disruptive to Tli structure and is therefore expected to cause a more profound phenotype. To explore this issue, we asked whether a Δ*tli* null strain phenocopies the transposon mutants on X-gluc indicator medium, and surprisingly found that Δ*tli* cells are not hyper-permeable (**Fig. 5A**). This result could reflect the acquisition of an inactivating mutation in *tle* during construction of the Δ*tli* strain. However, the hyper-permeable phenotype is induced when truncated Tli proteins lacking the N-terminal 23 (Tli-Δ23) and 51 (Tli-Δ51) residues are expressed from a plasmid in *Δtli* cells (**Fig. 5A**). Further truncation of the immunity protein to Tli-Δ75, which approximates the fragment produced by *tli-231* allele, does not support auto-permeabilization (**Fig. 5A**). These data suggest that Δ*tli* cells do not deploy active Tle. Accordingly, we found that *tle^+^ Δtli* cells behave like Δ*tssM* mock inhibitors when competed against *Δtle Δtli* target cells (**Fig. 5B**). The competitive fitness of the *tle^+^ Δtli* strain is nearly restored to wild-type levels when complemented with plasmid-borne *tli* (**Fig. 5B**). Furthermore, ECL *tle^+^ Δtli* cells still inhibit *E. coli* target bacteria (**Fig. 5C**), demonstrating that the Δ*tli* mutation has little effect on the delivery of other T6SS-1 effector proteins. Together, these results indicate Tle-mediated growth inhibition depends on a cytosolic pool of Tli.

**Fig. 5.**
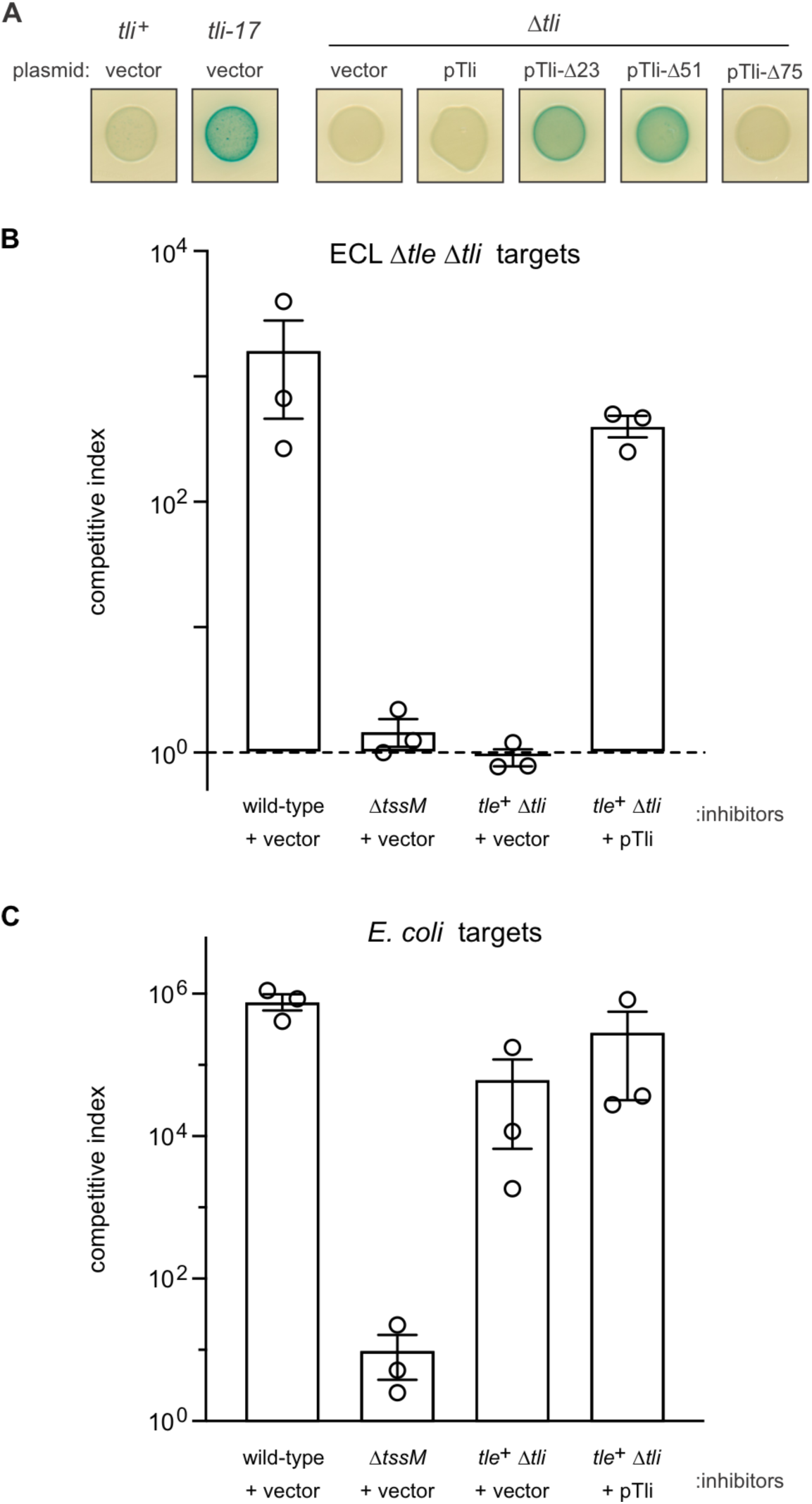
Tli is required to deploy Tle. **A**) ECL Δ*tli* null mutants do not exhibit hyper-permeability. The indicated strains were grown on LB-agar supplemented with X-gluc. Where indicated, strains carried an empty vector or were complemented with plasmids that produce Tli variants. **B**) The indicated ECL inhibitor strains were mixed with ECL Δ*tle Δtli* target cells and spotted onto LB-agar. After 4 h of co-culture, inhibitor and target cells were enumerated as colony forming units (cfu) on selective media. The competitive index equals the final ratio of inhibitor to target cells divided by the initial ratio. Presented data are the average ± standard error from three independent experiments. **C**) ECL inhibitor strains were mixed with *E. coli* X90 target cells and spotted onto LB-agar. Co-culture conditions and cfu enumerations were the same as described for panel **B**.

### Tli is required for the toxic activity of Tle

Tli could be required to load Tle into the T6SS-1 apparatus for export. We first tested whether Tli is necessary for effector secretion but were unable to detect Tle in culture supernatants from wild-type ECL cells. Therefore, we asked whether Tli is still required for growth inhibition activity when phospholipase delivery is ensured by fusion to the VgrG β-spike protein. We reasoned that the lipase domain could be fused to VgrG2 from ECL, because other *Enterobacter* strains encode ‘evolved’ VrgG2 homologs that carry effector domains at their C-termini (**Fig. S1**). AlphaFold2 modeling indicates that Tle is composed of two domains connected by a flexible linker (**Fig. 6A**). The novel N-terminal domain is presumably required to link the effector to VgrG for export, and the C-terminal domain adopts an α/β-hydrolase fold that resembles neutral lipases from fungi. Using the model as a guide, we fused the linker region and phospholipase domain of Tle to VgrG2 (**Fig. 6B**), then asked whether the chimera restores growth inhibition activity to a VgrG-deficient mutant. Because ECL Δ*tle-tli-vgrG1-vgrG2* cells do not produce functional VgrG, they cannot assemble the T6SS-1 apparatus (29) and do not kill *E. coli* target bacteria during co-culture (**Fig. 6C**). Complementation of Δ*tle-tli-vgrG1-vgrG2* cells with the VgrG2-lipase construct restores growth inhibition activity against *E. coli* target cells (**Fig. 6C**).

**Fig. 6.**
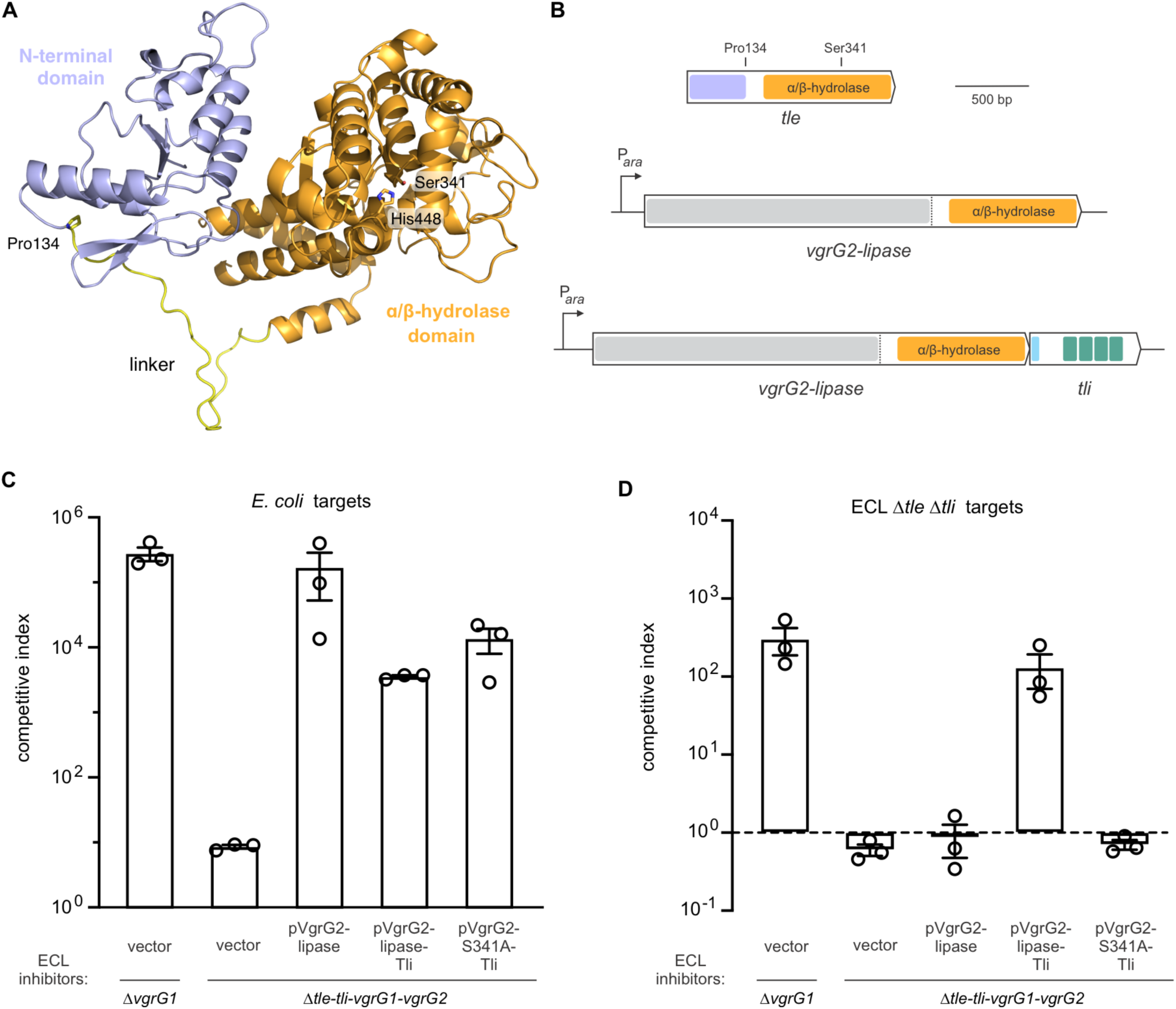
Tli is required to activate Tle prior to T6SS-mediate export. **A**) AlphaFold2 model of Tle. The C-terminal α/β-hydrolase domain is rendered in light orange and the linker region in yellow. The locations of predicted catalytic residues Ser341 and His448 are indicated. **B**) VgrG2-lipase domain fusion schematic. Coding sequence for the Tle linker region (beginning at Pro134) and lipase domain was fused to *vgrG2* under the control of an arabinose-inducible promoter. A matched construct that also includes the wild-type *tli* gene was also generated. **C**) The VgrG2-lipase fusion protein supports T6SS-1 activity. The indicated ECL inhibitor strains were mixed with *E. coli* X90 target cells and spotted onto LB-agar supplemented with L-arabinose. After 4 h of co-culture, inhibitor and target cells were enumerated as colony forming units (cfu) on selective media. The competitive index equals the final ratio of inhibitor to target cells divided by the initial ratio. Presented data are the average ± standard error from three independent experiments. **D**) ECL inhibitor strains were mixed with ECL Δ*tle Δtli* target cells and spotted onto LB-agar supplemented with L-arabinose. Co-culture conditions and cfu enumerations were the same as described for panel **C**.

Moreover, this inhibition activity is quantitatively similar that of an ECL Δ*vgrG1* inhibitor strain (**Fig. 6C**), which produces VgrG2 and Tle as separate proteins. These results suggest that the VgrG2-lipase chimera supports the same T6SS-1 activity as wild-type VgrG2. However, the VgrG2-lipase construct has no growth inhibition activity against ECL *Δtle Δtli* target cells (**Fig. 6D**), indicating that the fused lipase domain is inactive. We next tested whether addition of the *tli* gene to the fusion construct promotes phospholipase activity (**Fig. 6B**). Although the resulting VgrG2-lipase/Tli construct is somewhat less effective against *E. coli* target cells (**Fig. 6C**), it inhibits the growth of ECL *Δtle Δtli* target cells to the same extent as an ECL *ΔvgrG1* inhibitor strain (**Fig. 6D**). We also showed that mutation of the predicted catalytic Ser341 residue in the lipase domain abrogates growth inhibition activity against ECL Δ*tle Δtli* (**Fig. 6D**), but not *E. coli* target cells (**Fig. 6C**). Together, these results demonstrate that the Tle phospholipase domain is only active when co-produced with its cognate immunity protein.

## Discussion

Here, we identify ECL transposon mutants that exhibit increased permeability to a chromogenic β-glucuronidase substrate. The transposon insertions disrupt *tli*, which encodes an immunity protein that neutralizes the previously described Tle phospholipase effector (29). Thus, unmitigated Tle activity damages cellular membranes, allowing chromogenic substrate to enter the cytosol and/or β-glucuronidase to escape into the extracellular medium. This hyperpermeability is dependent on T6SS-1 activity, indicating that the mutants are intoxicated by Tle delivered from neighboring siblings rather than their own internally produced phospholipase. This phenomenon is similar to observations reported by Russell *et al.*, who found that deletion of the *tli5*^PA^ (PA3488) immunity gene from *Pseudomonas aeruginosa* PAO1 leads to increased cell permeability through unopposed PldD^PA^ phospholipase activity (30). However, in contrast to *P. aeruginosa*, ECL Δ*tli* null mutants fail to deploy active phospholipase and consequently do not exhibit hyperpermeability. Instead, the strongest phenotypes are associated with mutations that prevent targeting of Tli to the periplasm, yet still allow the immunity protein to accumulate to wild-type levels. These findings indicate that Tli has different functions depending on its subcellular localization. Tli in the periplasm acts to neutralize incoming phospholipase effectors, whereas cytoplasmic Tli is required for either the activation or export of Tle. Given that growth inhibition activity remains *tli*-dependent when phospholipase delivery is achieved through fusion to VgrG2, we propose that Tli activates the effector domain prior to T6SS-mediated export. In wild-type ECL, cytosolic Tli could be produced from alternative translation initiation codons found downstream of the signal sequence. The activation mechanism remains unknown, but cytoplasmic Tli could act as a binding chaperone to ensure proper folding of newly synthesized Tle. Because presumably only the effector protein is exported, this model suggests that there must be an additional mechanism to dissociate the Tle•Tli complex prior to loading into the T6SS apparatus.

Russell *et al.* first described and delineated five families of T6SS phospholipase effectors, which they designated Tle1 through Tle5 (30). Although not identified in that study, Tle^ECL^ from ECL contains the GxSxG catalytic motif found in the Tle1, Tle2, Tle3 and Tle4 effector families. AlphaFold2 modeling suggests that Tle^ECL^ adopts an α/β-hydrolase fold with the GxSxG sequence forming a classic nucleophilic ‘elbow’ in the turn between β4 and α7. A DALI server search indicates that Tle^ECL^ is not closely related to known T6SS effectors and instead is most similar to a family of neutral lipases secreted by fungi (31-33). This conclusion is supported by direct comparisons of the Tle^ECL^ model with the crystal structures of Tle1^PAO1^ and Tle4^PAO1^ from *Pseudomonas aeruginosa* PAO1. Superimposition of core α/β-hydrolase domains reveals that the flanking secondary elements are distinct between the effectors (**Fig. S2**). Moreover, Tle1^PAO1^ and Tle4^PAO1^ have modules that are absent from Tle^ECL^. Tle1^PAO1^ contains a putative membrane anchoring domain (34), and the α/β-hydrolase core of Tle4^PAO1^ is interrupted by three inserted segments that form a cap domain to cover the active site (35). We note that the Tle^ECL^ model also differs substantially from AlphaFold2 predictions for Tle2 (VasL) from *Vibrio cholerae* and Tle3 from *Burkholderia thailandensis* E264. Database searches reveal that Tle^ECL^ is commonly encoded by *Enterobacter* and *Cronobacter* T6SS loci, and it shares significant sequence identity with the Slp effector from *Serratia marcescens* (**Table S1**) (36). Homologous lipase domains are more broadly distributed across *Erwinia, Dickeya, Burkholderia* and *Vibrio* species, where they are fused to predicted VgrG, PAAR and DUF4150 domain proteins (**Table S1**). Related lipase domains also form the C-termini of rearrangement hotspot (Rhs) proteins in various *Pseudomonas* and *Vibrio* isolates. Rhs proteins constitute another important class of T6SS effectors that carry variable C-terminal toxin domains (1, 37-39). Finally, a handful of deltaproteobacteria encode predicted CdiA proteins with similar lipase domains (**Table S1**), indicating that this effector family can also be delivered through the T5SS pathway. Notably, these lipase sequences are linked to putative immunity genes that encode ankyrin-repeat proteins (**Table S1**), raising the possibility that all of these effectors are activated by their cognate immunity proteins.

**Table 1.**
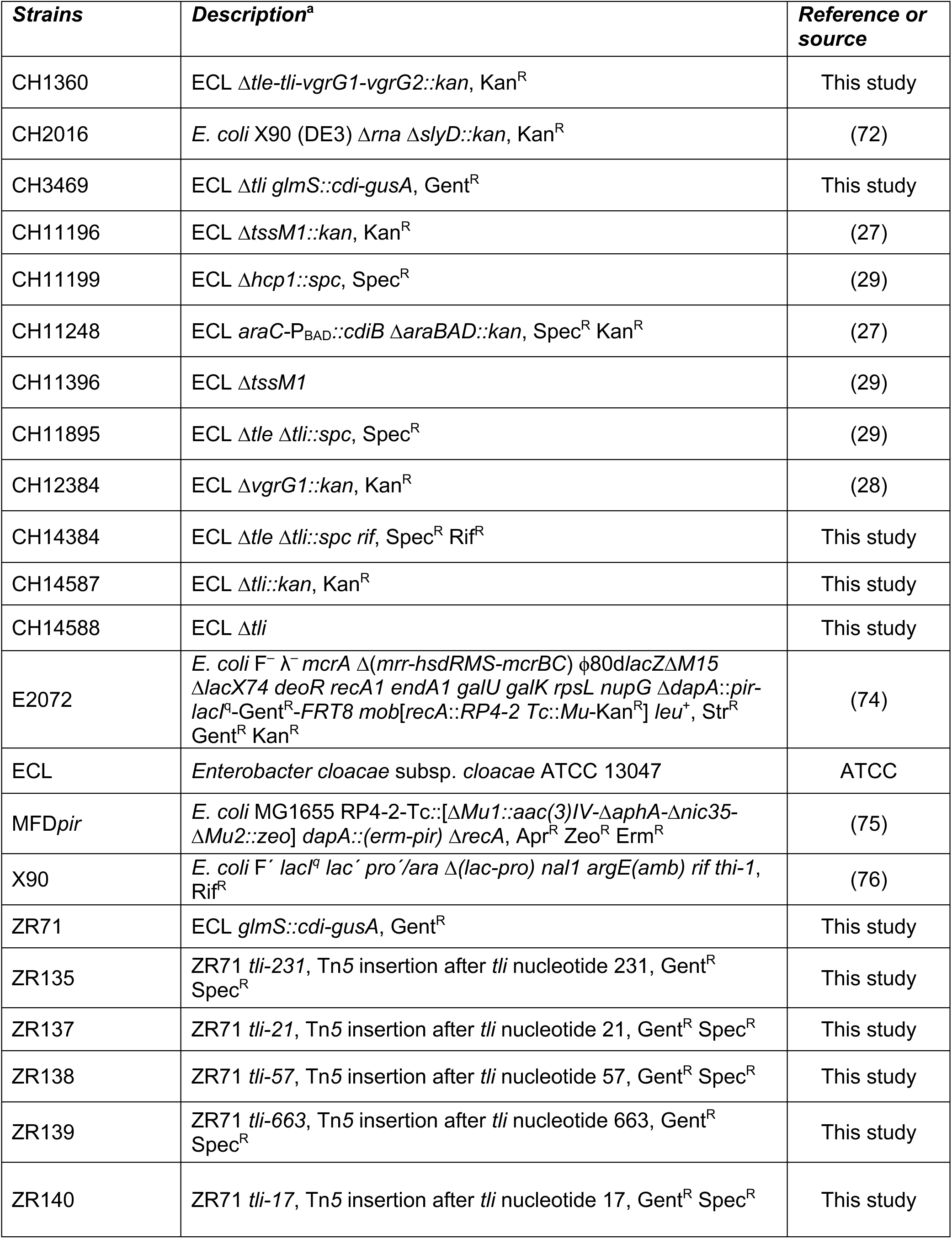
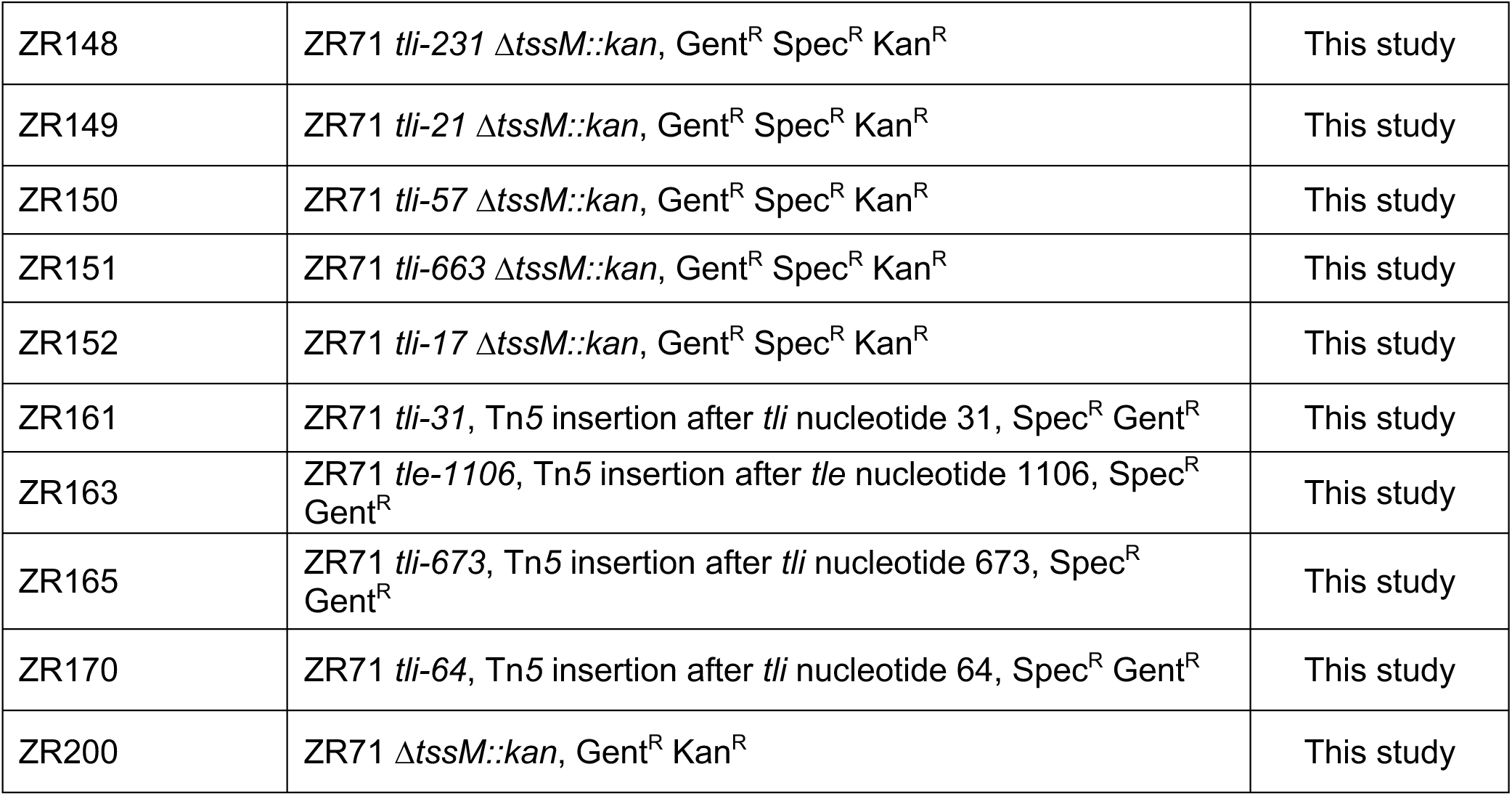
Bacterial strains

Though it remains to be determined whether other lipases require activation, a similar phenomenon has been reported for the PAAR domain-containing PefD effector of *Proteus mirabilis* HI4320. Mobley and coworkers found that disruption of the corresponding *pefE* immunity gene, which encodes a predicted β-propeller protein, abrogates the growth inhibition activity of PefD in co-cultures (40). PefD function is also dependent on *pefF* and *pefG*, which encode an armadillo repeat protein and a thioredoxin-like oxidoreductase, respectively (40, 41). AlphaFold2 indicates that PefD contains an intricate array of six disulfide bonds (https://www.uniprot.org/uniprotkb/B4EVA8/entry#structure), suggesting that PefE and PefF could be chaperones that hold the effector in the proper conformation for PefG-catalyzed disulfide bond formation. However, PefD and the related Tse7 effector from *P. aeruginosa* are toxic DNases that must transferred into the cytoplasm to inhibit growth (40, 42). Given that any disulfides would likely be reduced by glutathione and thioredoxin after delivery into the target cell, perhaps these bonds promote export rather than enzymatic activity. Proper disulfide formation is clearly important for other T6SS effectors that function in the periplasm. Peptidoglycan amidase and pore-forming effectors from *Serratia marcescens* both depend on DsbA (43), which is a highly conserved oxidoreductase that catalyzes disulfide formation in the periplasm of most Gram-negative bacteria (44). T5SS/CdiA effector domains are also known to exploit target-cell proteins to promote toxicity. CdiA from uropathogenic *E. coli* delivers a novel tRNA anticodon nuclease domain that is only active when bound to the cysteine biosynthetic enzyme CysK (26, 45, 46). Other CdiA proteins carry BECR-fold RNases that interact with elongation factor-Tu (EF-Tu) to cleave the 3’-acceptor stems of specific tRNA molecules (47-50).

Given that our screen ultimately selected for hyper-permeable cells, it is perhaps surprising that we did not recover other mutations that are known to disrupt the cell envelope. The first “leaky” mutants were isolated in screens for *E. coli* and *Salmonella* Typhimurium cells that release periplasmic RNase I and alkaline phosphatase into culture media (51-54). These early studies uncovered a critical role for TolA in Gram-negative cell envelope integrity (51). Later, *E. coli lamB* cells were used to identify mutations in *ompC* and *ompF* that allow high-molecular weight maltodextrins to enter the periplasm (55, 56). The same strategy also led to the discovery of *lptD* (57), which encodes an essential β-barrel protein that inserts lipopolysaccharide into the outer membrane (58-61). More recently, Bernhardt and co-workers used a cell-impermeable β-galactosidase substrate to identify over 100 *E. coli* genes that contribute to envelope integrity (62). Although the approach taken by Paradis-Bleu *et al.* is essentially the same as in our study, they note that disruption of outer-membrane integrity alone is probably sufficient for substrate entry in their screen (62). By contrast, ECL lacks outer- and inner-membrane transporters for β-glucuronides (63), and therefore both membranes must be permeabilized for substrate entry. These observations suggest that T6SS-delivered Tle damages the outer and inner membranes of target bacteria, and that the loss of Tli immunity function produces the most striking phenotype on indicator media.

## Materials and Methods

### Bacterial strain and plasmid constructions

Bacterial strains and plasmids are listed in **Tables 1** and **2** (respectively, and oligonucleotides are listed **Table S2**. All bacteria were grown at 37 °C in lysogeny broth (LB): 1% tryptone, 1% yeast extract, 0.5% NaCl. Media were supplemented with antibiotics at the following concentrations: ampicillin (Amp), 150 µg/mL; gentamicin (Gent), 12 μg/mL; kanamycin (Kan), 50 μg/mL; rifampicin (Rif), 200 µg/mL; spectinomycin (Spc), 15 μg/mL; tetracycline (Tet), 100 μg/mL and trimethoprim (Tp), 100 µg/mL, where indicated. The ECL *cdi-gusA* reporter strain ZR71 was constructed by Tn*7* mediated transposition. *E. coli gusA* was amplified with oligonucleotide primers CH3025/CH3026 and ligated to pUC18T-mini-Tn7T-Gm (64) using EcoRI/SacI restriction sites to generate plasmid pZR42. The ECL *cdiB* promoter region was amplified with CH3146/CH3147 and ligated to pZR42 using KpnI/EcoRI restriction sites to generate pCH3468. pCH3468 and pTNS2 (65) were introduced into wild-type ECL by conjugation and gentamicin-resistant (Gent^R^) clones were screened for *cdi-gusA* transposition into the *glmS* locus. The same mating procedure was used to generate strain CH3469 (ECL Δ*tli glmS::cdi-gusA*) from strain CH14588. The Δ*tssM1::kan* allele (from pCH11050) was introduced into ECL strains carrying plasmid pKOBEG using Red-mediated recombination as described (27, 29). ECL Δ*tli::kan* (CH14587) and Δ*tle-tli-vgrG1-vgrG2::kan* (CH1360) mutants were constructed using pCH371, which is a pRE118 (66) derivative that carries the *pheS(A294G)* counter-selectable marker instead of *sacB*. The *pheS(A294G)* allele was first amplified with DL2217/DL2194 and fused to the *tet* promoter in pBR322 using EcoRV/SphI restriction sites to generate plasmid pDAL912. The P*_tet_*-*pheS(A294G)* fragment was then amplified with DL2195/DL2196 and combined with pRE118-derived fragments generated with DL2197/DL2198 and D2199/DL1667 using overlap-extension PCR. The final PCR product was introduced into pRE118 using Red-mediated recombination to generate plasmid pDAL6480. A chloramphenicol-resistance (Cm^R^) cassette was amplified with CH5269/CH5270, digested with XbaI/NsiI and ligated to XbaI/SbfI-digested pDAL6480 to generate plasmid pCH377. The FRT-flanked Kan^R^ cassette from pKAN/pCH70 (67) was subcloned into pCH377 via SacI/KpnI restriction sites to generate pCH371. The pCH1359 Δ*tle-tli-vgrG1-vgrG2* knockout construct was generated by sequential cloning of CH3373/CH3374 (SacI/BamHI) and CH3197/CH3198 (XhoI/KpnI) fragments into pCH371. The pCH3884 Δ*tli::kan* knockout construct was generated by sequential cloning of ZR39/ZR40 (SacI/BamHI) and CH3377/CH3378 (XhoI/KpnI) fragments into pCH371. Plasmids pCH1359 and pCH3884 were introduced into ECL by conjugation followed by selection for Kan^R^ Cm^R^ exconjugants. Clones were cultured in M9 minimal medium supplemented 0.4% D-glucose, 2 mM D/L-chlorophenylalanine and 50 µg/mL kanamycin, then screened for Kan^R^ Cm^S^ clones that had lost the *pheS(A294G)* marker through homologous recombination. The Kan^R^ cassette was removed from CH14587 using the FLP recombinase encoded on pCP20 (68) to generate strain CH14588.

**Table 2.** Plasmids

| Plasmid | Description<sup>a</sup> | Reference |
| --- | --- | --- |
| pBR322 | cloning vector, Amp<sup>R</sup> Tet<sup>R</sup> | (77) |
| pCH70 | pBluescript with FRT-flanked kanamycin-resistance cassette ligated into SmaI restriction site, Amp<sup>R</sup> Kan<sup>R</sup> | (78) |
| pCH371 | pCATKAN-*pheS*(A294G), Cm<sup>R</sup> Kan<sup>R</sup> | This study |
| pCH377 | pCAT-*pheS*(A294G), Cm<sup>R</sup> | This study |
| pCH417 | pCH450::*vgrG2*(ECL)-CT/*vgrI*(Eh49162), Tet<sup>R</sup> | This study |
| pCH450 | pACYC184 derivative with *E. coli* *araC* and *araBAD* promoter for arabinose-inducible expression, Tet<sup>R</sup> | (67) |
| pCH494 | pCH450K::*tli*, Tet<sup>R</sup> | (29) |
| pCH495 | pCH450K::*tli*(Δ23), Tet<sup>R</sup> | This study |
| pCH1359 | pCATKAN-*pheS*(A294G)-Δ*tle-tli-vgrG1-vgrG2*, Cm<sup>R</sup> Kan<sup>R</sup> | This study |
| pCH3291 | pET21::*tli*(Δ23), Amp<sup>R</sup> | This study |
| pCH3468 | pUC18T-mini-Tn7T-Gm with the ECL *cdi* promoter region fused to *E. coli* *gusA* within the mini-Tn7, Amp<sup>R</sup> Gent<sup>R</sup> | This study |
| pCH3763 | pCH450::*vgrG2-lipase*(S341A)-*tli*, Tet<sup>R</sup> | This study |
| pCH3884 | pCATKAN-*pheS*(A294G)-Δ*tli*, Cm<sup>R</sup> Kan<sup>R</sup> | This study |
| pCH5036 | pCH450K::*tli*(Δ51), Tet<sup>R</sup> | This study |
| pCH5037 | pCH450K::*tli*(Δ75), Tet<sup>R</sup> | This study |
| pCH6118 | pET21b::*cdiB*(STEC3), Amp<sup>R</sup> | This study |
| pCH9062 | pCH450::*vgrG2-lipase-tli*, Tet<sup>R</sup> | This study |
| pCH9063 | pCH450::*vgrG2-lipase*, Tet<sup>R</sup> | This study |
| pCH9384 | pBluescript with FRT-flanked spectinomycin-resistance cassette ligated into SmaI restriction site, Amp<sup>R</sup> Spec<sup>R</sup> | (37) |
| pCH11050 | pCH70 with regions upstream and downstream of ECL *tssM1*, Amp<sup>R</sup> Kan<sup>R</sup> | (27) |
| pCH11198 | pCH450::*tssM*, Tet<sup>R</sup> | This study |
| pCP20 | temperature sensitive replication origin, expresses FLP recombinase. Amp<sup>R</sup>, Cm<sup>R</sup> | (79) |
| pDAL912 | pBR322::*pheS*(A294G), Amp<sup>R</sup> | This study |
| pDAL6480 | pRE118-*pheS*(A294G), Kan<sup>R</sup> | This study |
| pKOBEG | temperature sensitive replication origin, expresses the Bet-Gam-Exo proteins from phage $\lambda$. Cm<sup>R</sup> | (68) |
| pLG99 | Tn5 transposase expression vector harboring a mini-Tn5 derivative with P<sub>*rhaB*</sub>-out, Amp<sup>R</sup> Tmp<sup>R</sup> | (69) |
| pLG99-*spec* | pLG99 with a spectinomycin-resistance cassette cloned into Tn5, Amp<sup>R</sup> Spec<sup>R</sup> | This study |
| pRE118 | vector plasmid for *sacB* allelic exchange, Kan<sup>R</sup> | (66) |
| pSCBAD | pBBR1 derivative that carries *araC* and P<sub>BAD</sub> promoter, Tp<sup>R</sup> | (71) |
| pTNS2 | expresses Tn7 transposase, Amp<sup>R</sup> | (65) |
| pUC18T-mini-Tn7T-Gm | Tn7 transposase expression vector harboring mini-Tn7 with a gentamicin-resistance cassette, Amp<sup>R</sup> Gent<sup>R</sup> | (64) |
| pZR42 | pUC18T-mini-Tn7T-Gm with *E. coli gusA*, Amp<sup>R</sup> Gent<sup>R</sup> | This study |

The Tn*5* delivery vector pLG99 (69) was modified to introduce a spectinomycin-resistance (Spc^R^) cassette. An EcoRV/SpeI fragment from pSPM/pCH9384 (28) was ligated to a pLG99 vector prepared by NsiI (followed by end-filling with T4 DNA polymerase) and AvrII digestion. The ECL *tli* coding sequence was amplified with primer pairs, CH3418/CH3419 (wild-type *tli*), CH3719/CH3419 (*tli-Δ23*), CH5137/CH3419 (*tli-Δ51*) and CH5318/CH3419 (*tli-Δ75*), and the products ligated to pCH450K (67) using KpnI/XhoI restriction sites to generate plasmids pCH494, pCH495, pCH5036 and pCH5037, respectively. Primers CH3719/CH4154 were used amplify *tli(Δ23)* for ligation into pET21K via KpnI/XhoI to generate pCH3291, which over-produces cytosolic Tli with a C-terminal His_6_ epitope. ECL *tssM* was amplified with CH3023/CH3024 and ligated to pCH450 using NcoI/XhoI restriction sites to generate plasmid pCH11198. The *cdiB* gene from *E. coli* STEC_O31 was amplified with ZR443/ZR444 and ligated to pET21b using EcoRI/XhoI restriction sites to generate plasmid pCH6118.

To construct the VgrG2-lipase fusion, *vgrG2* from ECL was amplified with CH4452/CH5205 and ligated to pCH450 using EcoRI/KpnI restriction sites. *vgrG-CT/vgrI* sequences were amplified from *Enterobacter hormaechei* ATCC 49162 using CH5203/CH5204 and appended to the *vgrG2* construct using KpnI/PstI restriction sites to generate pCH417. Plasmid pCH417 was amplified with CH4452/CH5770 to add a SacI linker, and lipase domain/*tli* modules from CH5772/ZR248 and CH5772/CH3419 amplifications were ligated via SacI/XhoI to generate plasmids pCH9063 and pCH9062, respectively. The Ser341Ala substitution was introduced into the lipase domain by OE-PCR using primers CH4762/CH4763 in conjunction with CH5772/CH3419 to generate plasmid pCH3763.

### Transposon mutagenesis and mutant screening

Plasmid pLG99::*spec* was introduced into ECL strain ZR71 cells via conjugation and Spc^R^ clones selected on tryptone broth-agar (1% tryptone, 0.5% NaCl, 1.5% agar) supplemented with 100 µg/mL Spc and 100 μg/mL of 5-bromo-4-chloro-1H-indol-3-yl β-D-glucopyranosiduronic acid (X-gluc). Transposon insertion sites were identified by arbitrarily-primed PCR. The Tn*5* right arm was amplified with Tn5-Right1/CH2020 or Tn5-Right1/CH2022. Products were re-amplified using nested primers Tn5-Right2/CH2021. The Tn*5* left arm was amplified using primers Tn5-Left1/CH2020 and Tn5-Left1/CH2022. The resulting reactions were amplified with nested primers Tn5-Left2/CH2021. Second-round reactions were sequenced using oligonucleotides Tn5-Right3 and Tn5-Left3 as primers.

### X-gluc permeability and β-glucuronidase assays

β-glucuronidase activity was quantified as described by Miller (70). Cells were harvested from tryptone broth-agar and suspended in 100 mM sodium phosphate (pH 7.0) at an optical density at 600 nm (OD_600_) of 1.0. A drop of toluene and a drop of 0.1% SDS was added to 100 µL of the cell suspension followed by vortexing for 15 s. The suspension was then incubated at 37 °C for 90 min without caps to allow evaporation of toluene. Cell suspensions and substrate solution (1 mg/mL *p*-nitrophenyl β-D-glucuronide in 100 mM sodium phosphate (pH 7.0), 5 mM β-mercaptoethanol) were equilibrated to 28 °C. Cell lysate (100 μL) was added to 600 μL of substrate solution and incubated at 28 °C for 30 min. Reactions were quenched with 700 µL of 1 M sodium carbonate. Quenched reactions were centrifuged at 14,000 5*g* for 10 min and supernatant was removed and absorbance measured at 420 nm. β-glucuronidase activity was quantified in Miller units, where enzyme units = 1,000 (A_420_)/(1.0 OD_600_)(0.1 mL)(30 min). Overnight cultures were adjusted to OD_600_ = 3.0 and 10 μL was spotted onto TB-agar supplemented with 100 μg/mL X-gluc and incubated overnight at 37 °C. Culture plates were imaged using a light scanner.

### Competition co-cultures

Inhibitor and target cells were grown overnight in LB, then diluted 1:50 into fresh LB and grown at 37 °C to mid-log phase. Cells were harvested by centrifugation at 3,400 x f for 2 min, resuspended to an optical density at 600 nm (OD_600_) of 3.0 and mixed at a 10:1 ratio of inhibitor to target cells. Cell mixtures (10 µL) were spotted onto LB-agar (supplemented with 0.4% L-arabinose when using strains with arabinose-inducible constructs) and incubated at 37 °C. A portion of each cell mixture was serially diluted into 15 M9 salts and plated onto antibiotic supplemented LB-agar to enumerate inhibitor and target cells as colony forming units (cfu) at *t* = 0 h. For the competition in Fig. 4B, inhibitor strains were scored using Tp^R^ (conferred by plasmid pSCBAD (71)) and targets were scored by Spc^R^. For all other competitions, inhibitors were scored by Tet^R^ (conferred by pCH450 derivatives), and *E. coli* and ECL target cells were enumerated by Rif^R^. After 4 h of co-culture, cells were harvested from the agar surface with polyester-tipped swabs and resuspended in 800 µL of 1 x M9 salts. The cell suspensions were serially diluted in 1 x M9 salts and plated as described above to quantify end-point cfu. Competitive indices were calculated as the final ratio of inhibitor to target cells divided by the initial ratio of inhibitor to target cells.

### Hcp secretion

Overnight cultures of ECL were diluted 1:50 into 3 mL LB and grown to OD_600_ ∼ 1.0 at 37 °C. The cultures were then supplemented to 0.4% L-arabinose and incubated for 45 min. The cultures were centrifuged at 3,400 x f for 5 min, and the cell pellets resuspended in 100 µL of urea lysis buffer [8 M urea, 150 mM NaCl, 20 mM Tris-HCl (pH 8.0)] for lysis. Supernatants were collected and centrifuged for an additional 5 min to remove any remaining cells. Supernatants were then passed through a 0.22 µm low-binding polyvinylidene fluoride (PVDF) filter. The filtered culture supernatants were adjusted to 10% trichloroacetic acid and incubated on ice for 1 h. Samples were centrifuged at 21,000 x for 10 min. Precipitates were resuspended twice in 1 mL ice-cold acetone and recollected by centrifugation at 21,000 x for 5 min. Precipitated were air-dried at ambient temperature then dissolved in 50 µL urea lysis buffer. Protein concentrations were estimated by Bradford assay and ∼5 µg loaded onto 10% polyacrylamide SDS gels for electrophoresis and immunoblot analysis.

### Antisera generation and immunoblotting

His_6_-tagged variants of Tli(Δ23) from ECL and CdiB from *E. coli* STEC_O31 were over-produced in *E. coli* strain CH2016 from plasmids pCH3291 and pCH6118, respectively. Overnight cultures were diluted 1:100 in fresh LB supplemented with Amp and grown to mid-log phase at 37°C in a baffled flask. Expression was then induced with isopropyl β-D-1-thiogalactopyranoside at a final concentration of 1.5 mM for 90 min. Cultures were harvested by centrifugation and cell pellets frozen at −80°C. Cells were broken by freeze–thaw in urea lysis buffer supplemented with 0.05% Triton X-100 and 20 mM imidazole as described previously (72). Proteins were purified by Ni^2+^-affinity chromatography under denaturing conditions in urea lysis buffer, followed by elution in urea lysis buffer supplemented with 25 mM of ethylenediaminetetraacetic acid. The purified proteins were dialyzed against water and lyophilized to be used as antigens to generate rabbit polyclonal antisera (Cocalico Biologicals Inc., Reamstown, PA).

Samples were analyzed by SDS-polyacrylamide gel electrophoresis using Tris-tricine buffered gels run at 110 V. For Hcp and CdiB immunoblots, samples were run on 10% polyacrylamide gels for 1 h. Tli samples were run on 10% polyacrylamide gels for 2 h 15 min, and CdiA samples were run on 6% polyacrylamide gels for 3.5 h. Gels were soaked for 10 min in transfer buffer [25 mM Tris, 192 mM glycine (pH 8.6), 20% methanol (10% methanol for CdiA gels)] before electroblotting to PVDF membranes using a semi-dry transfer apparatus run at 17 V for 30 min. For CdiA, gels were electroblotted for 1 h. Membranes were blocked with 4% non-fat milk in 1 x PBS for 45 min at ambient temperature, then incubated with primary antibodies in 1 x PBS, 4% non-fat milk overnight. Rabbit polyclonal antisera for Hcp (29), Tli, CdiA (73) and CdiB were used at dilutions of 1:10,000, 1:5,000, 1:10,000, and 1:12,500, respectively. After three 10 min washes 1 x PBS, the membranes were incubated with IRDye 800CW-conjugated goat anti-rabbit IgG (LI-COR, 1:125,000 dilution) in 1 x PBS for 45 min. Immunoblots were visualized using a LI-COR Odyssey infrared imager.

## Author contributions

Conceptualization: S. J. J., Z. C. R., D. A. L. & C. S. H., Funding acquisition: D. A. L. & C. S. H.;

Investigation: S. J. J., Z. C. R., A. F. W., D. Q. N. & F. G. S.; Methodology: Z. C. R. & C. S. H.;

Supervision: D. A. L. & C. S. H.; Validation: S. J. J., D. Q. N. & F. G. S.; Visualization: Z. C. R. & C. S. H.; Writing – original draft preparation: Z. C. R. & C. S. H.; Writing – review and editing: S. J. J., D. A. L. & C. S. H.

## Acknowledgments

We thank Stephanie Aoki for the construction of plasmid pDAL6480 and Jiang Xiong for technical assistance. This work was supported by grants GM117930 (C.S.H.) from the National Institutes of Health and MCB 1545720 (D.A.L & C.S.H.) from the National Science Foundation. Z.C.R. was supported by the Tri-Counties Blood Bank Postdoctoral Fellowship.

## References

1. Ruhe ZC, Low DA, Hayes CS. 2020. Polymorphic Toxins and Their Immunity Proteins: Diversity, Evolution, and Mechanisms of Delivery. Annu Rev Microbiol 74:497–520.

2. Aoki SK, Pamma R, Hernday AD, Bickham JE, Braaten BA, Low DA. 2005. Contact-dependent inhibition of growth in *Escherichia coli*. Science 309:1245–8.

3. Ruhe ZC, Subramanian P, Song K, Nguyen JY, Stevens TA, Low DA, Jensen GJ, Hayes CS. 2018. Programmed Secretion Arrest and Receptor-Triggered Toxin Export during Antibacterial Contact-Dependent Growth Inhibition. Cell 175:921–933 e14.

4. Hood RD, Singh P, Hsu F, Guvener T, Carl MA, Trinidad RR, Silverman JM, Ohlson BB, Hicks KG, Plemel RL, Li M, Schwarz S, Wang WY, Merz AJ, Goodlett DR, Mougous JD. 2010. A type VI secretion system of *Pseudomonas aeruginosa* targets a toxin to bacteria. Cell Host Microbe 7:25–37.

5. MacIntyre DL, Miyata ST, Kitaoka M, Pukatzki S. 2010. The *Vibrio cholerae* type VI secretion system displays antimicrobial properties. Proc Natl Acad Sci U S A 107:19520–4.

6. Leiman PG, Basler M, Ramagopal UA, Bonanno JB, Sauder JM, Pukatzki S, Burley SK, Almo SC, Mekalanos JJ. 2009. Type VI secretion apparatus and phage tail-associated protein complexes share a common evolutionary origin. Proc Natl Acad Sci U S A 106:4154–9.

7. Brunet YR, Zoued A, Boyer F, Douzi B, Cascales E. 2015. The Type VI Secretion TssEFGK-VgrG Phage-Like Baseplate Is Recruited to the TssJLM Membrane Complex via Multiple Contacts and Serves As Assembly Platform for Tail Tube/Sheath Polymerization. PLoS Genet 11:e1005545.

8. Pukatzki S, Ma AT, Revel AT, Sturtevant D, Mekalanos JJ. 2007. Type VI secretion system translocates a phage tail spike-like protein into target cells where it cross-links actin. Proc Natl Acad Sci U S A 104:15508–13.

9. Shneider MM, Buth SA, Ho BT, Basler M, Mekalanos JJ, Leiman PG. 2013. PAAR-repeat proteins sharpen and diversify the type VI secretion system spike. Nature 500:350–3.

10. Silverman JM, Agnello DM, Zheng H, Andrews BT, Li M, Catalano CE, Gonen T, Mougous JD. 2013. Haemolysin coregulated protein is an exported receptor and chaperone of type VI secretion substrates. Mol Cell 51:584–93.

11. Hernandez RE, Gallegos-Monterrosa R, Coulthurst SJ. 2020. Type VI secretion system effector proteins: Effective weapons for bacterial competitiveness. Cell Microbiol 22:e13241.

12. Brunet YR, Khodr A, Logger L, Aussel L, Mignot T, Rimsky S, Cascales E. 2015. H-NS Silencing of the Salmonella Pathogenicity Island 6-Encoded Type VI Secretion System Limits Salmonella enterica Serovar Typhimurium Interbacterial Killing. Infect Immun 83:2738–50.

13. Caro F, Caro JA, Place NM, Mekalanos JJ. 2020. Transcriptional Silencing by TsrA in the Evolution of Pathogenic Vibrio cholerae Biotypes. mBio 11.

14. Storey D, McNally A, Åstrand M, Sa-Pessoa Graca Santos J, Rodriguez-Escudero I, Elmore B, Palacios L, Marshall H, Hobley L, Molina M, Cid VJ, Salminen TA, Bengoechea JA. 2020. Klebsiella pneumoniae type VI secretion system-mediated microbial competition is PhoPQ controlled and reactive oxygen species dependent. PLoS Pathog 16:e1007969.

15. Ma J, Sun M, Pan Z, Song W, Lu C, Yao H. 2018. Three Hcp homologs with divergent extended loop regions exhibit different functions in avian pathogenic Escherichia coli. Emerg Microbes Infect 7:49.

16. Bernard CS, Brunet YR, Gueguen E, Cascales E. 2010. Nooks and crannies in type VI secretion regulation. J Bacteriol 192:3850–60.

17. Pukatzki S, Ma AT, Sturtevant D, Krastins B, Sarracino D, Nelson WC, Heidelberg JF, Mekalanos JJ. 2006. Identification of a conserved bacterial protein secretion system in *Vibrio cholerae* using the *Dictyostelium* host model system. Proc Natl Acad Sci U S A 103:1528–33.

18. Bernard CS, Brunet YR, Gavioli M, Lloubès R, Cascales E. 2011. Regulation of type VI secretion gene clusters by sigma54 and cognate enhancer binding proteins. J Bacteriol 193:2158–67.

19. Sana TG, Hachani A, Bucior I, Soscia C, Garvis S, Termine E, Engel J, Filloux A, Bleves S. 2012. The second type VI secretion system of Pseudomonas aeruginosa strain PAO1 is regulated by quorum sensing and Fur and modulates internalization in epithelial cells. J Biol Chem 287:27095–105.

20. Majerczyk C, Schneider E, Greenberg EP. 2016. Quorum sensing control of Type VI secretion factors restricts the proliferation of quorum-sensing mutants. Elife 5.

21. Mougous JD, Gifford CA, Ramsdell TL, Mekalanos JJ. 2007. Threonine phosphorylation post-translationally regulates protein secretion in Pseudomonas aeruginosa. Nat Cell Biol 9:797–803.

22. Wäneskog M, Halvorsen T, Filek K, Xu F, Hammarlöf DL, Hayes CS, Braaten BA, Low DA, Poole SJ, Koskiniemi S. 2021. Escherichia coli EC93 deploys two plasmid-encoded class I contact-dependent growth inhibition systems for antagonistic bacterial interactions. Microb Genom 7.

23. Majerczyk C, Brittnacher M, Jacobs M, Armour CD, Radey M, Schneider E, Phattarasokul S, Bunt R, Greenberg EP. 2014. Global analysis of the Burkholderia thailandensis quorum sensing-controlled regulon. J Bacteriol 196:1412–24.

24. Rojas CM, Ham JH, Deng WL, Doyle JJ, Collmer A. 2002. HecA, a member of a class of adhesins produced by diverse pathogenic bacteria, contributes to the attachment, aggregation, epidermal cell killing, and virulence phenotypes of *Erwinia chrysanthemi* EC16 on *Nicotiana clevelandii* seedlings. Proc Natl Acad Sci U S A 99:13142–7.

25. Rojas CM, Ham JH, Schechter LM, Kim JF, Beer SV, Collmer A. 2004. The *Erwinia chrysanthemi* EC16 *hrp/hrc* gene cluster encodes an active Hrp type III secretion system that is flanked by virulence genes functionally unrelated to the Hrp system. Mol Plant Microbe Interact 17:644–53.

26. Aoki SK, Diner EJ, de Roodenbeke CT, Burgess BR, Poole SJ, Braaten BA, Jones AM, Webb JS, Hayes CS, Cotter PA, Low DA. 2010. A widespread family of polymorphic contact-dependent toxin delivery systems in bacteria. Nature 468:439–42.

27. Beck CM, Morse RP, Cunningham DA, Iniguez A, Low DA, Goulding CW, Hayes CS. 2014. CdiA from *Enterobacter cloacae* delivers a toxic ribosomal RNase into target bacteria. Structure 22:707–718.

28. Whitney JC, Beck CM, Goo YA, Russell AB, Harding BN, De Leon JA, Cunningham DA, Tran BQ, Low DA, Goodlett DR, Hayes CS, Mougous JD. 2014. Genetically distinct pathways guide effector export through the type VI secretion system. Mol Microbiol 92:529–42.

29. Donato SL, Beck CM, Ruhe ZC, Cunningham DA, Singleton I, Low DA, Hayes CS. 2020. The beta-encapsulation cage of rearrangement hotspot (Rhs) effectors is required for type VI secretion. Proc Natl Acad Sci U S A:in revision.

30. Russell AB, LeRoux M, Hathazi K, Agnello DM, Ishikawa T, Wiggins PA, Wai SN, Mougous JD. 2013. Diverse type VI secretion phospholipases are functionally plastic antibacterial effectors. Nature 496:508–12.

31. Brady L, Brzozowski AM, Derewenda ZS, Dodson E, Dodson G, Tolley S, Turkenburg JP, Christiansen L, Huge-Jensen B, Norskov L, et al. 1990. A serine protease triad forms the catalytic centre of a triacylglycerol lipase. Nature 343:767–70.

32. Lan D, Xu H, Xu J, Dubin G, Liu J, Iqbal Khan F, Wang Y. 2017. Malassezia globosa MgMDL2 lipase: Crystal structure and rational modification of substrate specificity. Biochem Biophys Res Commun 488:259–265.

33. Lou Z, Li M, Sun Y, Liu Y, Liu Z, Wu W, Rao Z. 2010. Crystal structure of a secreted lipase from Gibberella zeae reveals a novel “double-lock” mechanism. Protein Cell 1:760–70.

34. Hu H, Zhang H, Gao Z, Wang D, Liu G, Xu J, Lan K, Dong Y. 2014. Structure of the type VI secretion phospholipase effector Tle1 provides insight into its hydrolysis and membrane targeting. Acta Crystallogr D Biol Crystallogr 70:2175–85.

35. Lu D, Zheng Y, Liao N, Wei L, Xu B, Liu X, Liu J. 2014. The structural basis of the Tle4-Tli4 complex reveals the self-protection mechanism of H2-T6SS in Pseudomonas aeruginosa. Acta Crystallogr D Biol Crystallogr 70:3233–43.

36. Cianfanelli FR, Alcoforado Diniz J, Guo M, De Cesare V, Trost M, Coulthurst SJ. 2016. VgrG and PAAR Proteins Define Distinct Versions of a Functional Type VI Secretion System. PLoS Pathog 12:e1005735.

37. Koskiniemi S, Lamoureux JG, Nikolakakis KC, t’Kint de Roodenbeke C, Kaplan MD, Low DA, Hayes CS. 2013. Rhs proteins from diverse bacteria mediate intercellular competition. Proc Natl Acad Sci U S A 110:7032-7.

38. Alcoforado Diniz J, Coulthurst SJ. 2015. Intraspecies Competition in *Serratia marcescens* Is Mediated by Type VI-Secreted Rhs Effectors and a Conserved Effector-Associated Accessory Protein. J Bacteriol 197:2350–60.

39. Zhang D, de Souza RF, Anantharaman V, Iyer LM, Aravind L. 2012. Polymorphic toxin systems: Comprehensive characterization of trafficking modes, processing, mechanisms of action, immunity and ecology using comparative genomics. Biol Direct 7:18.

40. Alteri CJ, Himpsl SD, Zhu K, Hershey HL, Musili N, Miller JE, Mobley HLT. 2017. Subtle variation within conserved effector operon gene products contributes to T6SS-mediated killing and immunity. PLoS Pathog 13:e1006729.

41. Alteri CJ, Himpsl SD, Pickens SR, Lindner JR, Zora JS, Miller JE, Arno PD, Straight SW, Mobley HL. 2013. Multicellular bacteria deploy the type VI secretion system to preemptively strike neighboring cells. PLoS Pathog 9:e1003608.

42. Pissaridou P, Allsopp LP, Wettstadt S, Howard SA, Mavridou DAI, Filloux A. 2018. The Pseudomonas aeruginosa T6SS-VgrG1b spike is topped by a PAAR protein eliciting DNA damage to bacterial competitors. Proc Natl Acad Sci U S A 115:12519–12524.

43. Mariano G, Monlezun L, Coulthurst SJ. 2018. Dual Role for DsbA in Attacking and Targeted Bacterial Cells during Type VI Secretion System-Mediated Competition. Cell Rep 22:774–785.

44. Collet JF, Cho SH, Iorga BI, Goemans CV. 2020. How the assembly and protection of the bacterial cell envelope depend on cysteine residues. J Biol Chem 295:11984–11994.

45. Diner EJ, Beck CM, Webb JS, Low DA, Hayes CS. 2012. Identification of a target cell permissive factor required for contact-dependent growth inhibition (CDI). Genes Dev 26:515–25.

46. Johnson PM, Beck CM, Morse RP, Garza-Sánchez F, Low DA, Hayes CS, Goulding CW. 2016. Unraveling the essential role of CysK in CDI toxin activation. Proc Natl Acad Sci U S A 113:9792–7.

47. Jones AM, Garza-Sánchez F, So J, Hayes CS, Low DA. 2017. Activation of contact-dependent antibacterial tRNase toxins by translation elongation factors. Proc Natl Acad Sci U S A 114:E1951–e1957.

48. Michalska K, Gucinski GC, Garza-Sánchez F, Johnson PM, Stols LM, Eschenfeldt WH, Babnigg G, Low DA, Goulding CW, Joachimiak A, Hayes CS. 2017. Structure of a novel antibacterial toxin that exploits elongation factor Tu to cleave specific transfer RNAs. Nucleic Acids Res 45:10306–10320.

49. Gucinski GC, Michalska K, Garza-Sánchez F, Eschenfeldt WH, Stols L, Nguyen JY, Goulding CW, Joachimiak A, Hayes CS. 2019. Convergent Evolution of the Barnase/EndoU/Colicin/RelE (BECR) Fold in Antibacterial tRNase Toxins. Structure 27:1660–1674.e5.

50. Wang J, Yashiro Y, Sakaguchi Y, Suzuki T, Tomita K. 2022. Mechanistic insights into tRNA cleavage by a contact-dependent growth inhibitor protein and translation factors. Nucleic Acids Res 50:4713–4731.

51. Amouroux C, Lazzaroni JC, Portalier R. 1991. Isolation and characterization of extragenic suppressor mutants of the tolA-876 periplasmic-leaky allele in Escherichia coli K-12. FEMS Microbiol Lett 62:305–13.

52. Lazzaroni JC, Portalier RC. 1981. Genetic and biochemical characterization of periplasmic-leaky mutants of Escherichia coli K-12. J Bacteriol 145:1351–8.

53. Lopes J, Gottfried S, Rothfield L. 1972. Leakage of periplasmic enzymes by mutants of Escherichia coli and Salmonella typhimurium: isolation of “periplasmic leaky” mutants. J Bacteriol 109:520–5.

54. Weigand RA, Rothfield LI. 1976. Genetic and physiological classification of periplasmic-leaky mutants of Salmonella typhimurium. J Bacteriol 125:340–5.

55. Benson SA, Decloux A. 1985. Isolation and characterization of outer membrane permeability mutants in Escherichia coli K-12. J Bacteriol 161:361–7.

56. Benson SA, Occi JL, Sampson BA. 1988. Mutations that alter the pore function of the OmpF porin of Escherichia coli K12. J Mol Biol 203:961–70.

57. Sampson BA, Misra R, Benson SA. 1989. Identification and characterization of a new gene of Escherichia coli K-12 involved in outer membrane permeability. Genetics 122:491–501.

58. Dong H, Xiang Q, Gu Y, Wang Z, Paterson NG, Stansfeld PJ, He C, Zhang Y, Wang W, Dong C. 2014. Structural basis for outer membrane lipopolysaccharide insertion. Nature 511:52–6.

59. Gu Y, Stansfeld PJ, Zeng Y, Dong H, Wang W, Dong C. 2015. Lipopolysaccharide is inserted into the outer membrane through an intramembrane hole, a lumen gate, and the lateral opening of LptD. Structure 23:496–504.

60. Li X, Gu Y, Dong H, Wang W, Dong C. 2015. Trapped lipopolysaccharide and LptD intermediates reveal lipopolysaccharide translocation steps across the Escherichia coli outer membrane. Sci Rep 5:11883.

61. Qiao S, Luo Q, Zhao Y, Zhang XC, Huang Y. 2014. Structural basis for lipopolysaccharide insertion in the bacterial outer membrane. Nature 511:108–11.

62. Paradis-Bleau C, Kritikos G, Orlova K, Typas A, Bernhardt TG. 2014. A genome-wide screen for bacterial envelope biogenesis mutants identifies a novel factor involved in cell wall precursor metabolism. PLoS Genet 10:e1004056.

63. Liang WJ, Wilson KJ, Xie H, Knol J, Suzuki S, Rutherford NG, Henderson PJ, Jefferson RA. 2005. The gusBC genes of Escherichia coli encode a glucuronide transport system. J Bacteriol 187:2377–85.

64. Choi KH, Mima T, Casart Y, Rholl D, Kumar A, Beacham IR, Schweizer HP. 2008. Genetic tools for select-agent-compliant manipulation of Burkholderia pseudomallei. Appl Environ Microbiol 74:1064–75.

65. Choi KH, Gaynor JB, White KG, Lopez C, Bosio CM, Karkhoff-Schweizer RR, Schweizer HP. 2005. A Tn7-based broad-range bacterial cloning and expression system. Nat Methods 2:443–8.

66. Edwards RA, Keller LH, Schifferli DM. 1998. Improved allelic exchange vectors and their use to analyze 987P fimbria gene expression. Gene 207:149–57.

67. Hayes CS, Sauer RT. 2003. Cleavage of the A site mRNA codon during ribosome pausing provides a mechanism for translational quality control. Mol Cell 12:903–11.

68. Cherepanov PP, Wackernagel W. 1995. Gene disruption in *Escherichia coli*: TcR and KmR cassettes with the option of Flp-catalyzed excision of the antibiotic-resistance determinant. Gene 158:9–14.

69. Gallagher LA, Ramage E, Patrapuvich R, Weiss E, Brittnacher M, Manoil C. 2013. Sequence-defined transposon mutant library of Burkholderia thailandensis. mBio 4:e00604–13.

70. Miller JH. 1972. Experiments in Molecular Genetics. Cold Spring Harbor Laboratory Press, Cold Spring Harbor, NY.

71. Koskiniemi S, Garza-Sánchez F, Edman N, Chaudhuri S, Poole SJ, Manoil C, Hayes CS, Low DA. 2015. Genetic analysis of the CDI pathway from Burkholderia pseudomallei 1026b. PLoS One 10:e0120265.

72. Garza-Sánchez F, Janssen BD, Hayes CS. 2006. Prolyl-tRNA(Pro) in the A-site of SecM-arrested ribosomes inhibits the recruitment of transfer-messenger RNA. J Biol Chem 281:34258–68.

73. Ruhe ZC, Townsley L, Wallace AB, King A, Van der Woude MW, Low DA, Yildiz FH, Hayes CS. 2015. CdiA promotes receptor-independent intercellular adhesion. Mol Microbiol 98:175–92.

74. Zarzycki-Siek J, Norris MH, Kang Y, Sun Z, Bluhm AP, McMillan IA, Hoang TT. 2013. Elucidating the Pseudomonas aeruginosa fatty acid degradation pathway: identification of additional fatty acyl-CoA synthetase homologues. PLoS One 8:e64554.

75. Ferrières L, Hémery G, Nham T, Guérout AM, Mazel D, Beloin C, Ghigo JM. 2010. Silent mischief: bacteriophage Mu insertions contaminate products of Escherichia coli random mutagenesis performed using suicidal transposon delivery plasmids mobilized by broad-host-range RP4 conjugative machinery. J Bacteriol 192:6418–27.

76. Beckwith JR, Signer ER. 1966. Transposition of the lac region of Escherichia coli. I. Inversion of the lac operon and transduction of lac by phi80. J Mol Biol 19:254-65.

77. Bolivar F, Rodriguez RL, Greene PJ, Betlach MC, Heyneker HL, Boyer HW, Crosa JH, Falkow S. 1977. Construction and characterization of new cloning vehicles. II. A multipurpose cloning system. Gene 2:95–113.

78. Hayes CS, Bose B, Sauer RT. 2002. Proline residues at the C terminus of nascent chains induce SsrA tagging during translation termination. J Biol Chem 277:33825–32.

79. Pérez A, Canle D, Latasa C, Poza M, Beceiro A, Tomás Mdel M, Fernández A, Mallo S, Pérez S, Molina F, Villanueva R, Lasa I, Bou G. 2007. Cloning, nucleotide sequencing, and analysis of the AcrAB-TolC efflux pump of Enterobacter cloacae and determination of its involvement in antibiotic resistance in a clinical isolate. Antimicrob Agents Chemother 51:3247–53.

